# EslB is required for cell wall biosynthesis and modification in *Listeria monocytogenes*

**DOI:** 10.1101/2020.02.03.932061

**Authors:** Jeanine Rismondo, Lisa M. Schulz, Maria Yacoub, Ashima Wadhawan, Michael Hoppert, Marc S. Dionne, Angelika Gründling

## Abstract

Lysozyme is an important component of the innate immune system. It functions by hydrolysing the peptidoglycan (PG) layer of bacteria. The human pathogen *Listeria monocytogenes* is intrinsically lysozyme resistant. The peptidoglycan *N*-deacetylase PgdA and *O*-acetyltransferase OatA are two known factors contributing to its lysozyme resistance. Furthermore, it was shown that the absence of components of an ABC transporter, here referred to as EslABC, leads to reduced lysozyme resistance. How its activity is linked to lysozyme resistance is still unknown. To investigate this further, a strain with a deletion in *eslB*, coding for a membrane component of the ABC transporter, was constructed in *L. monocytogenes* strain 10403S. The *eslB* mutant showed a 40-fold reduction in the minimal inhibitory concentration to lysozyme. Analysis of the PG structure revealed that the *eslB* mutant produced PG with reduced levels of *O*-acetylation. Using growth and autolysis assays, we show that the absence of EslB manifests in a growth defect in media containing high concentrations of sugars and increased endogenous cell lysis. A thinner PG layer produced by the *eslB* mutant under these growth conditions might explain these phenotypes. Furthermore, the *eslB* mutant had a noticeable cell division defect and formed elongated cells. Microscopy analysis revealed that an early cell division protein still localized in the *eslB* mutant indicating that a downstream process is perturbed. Based on our results, we hypothesize that EslB affects the biosynthesis and modification of the cell wall in *L. monocytogenes* and is thus important for the maintenance of cell wall integrity.

**IMPORTANCE:** The ABC transporter EslABC is associated with the intrinsic lysozyme resistance of *Listeria monocytogenes*. However, the exact role of the transporter in this process and in the physiology of *L. monocytogenes* is unknown. Using different assays to characterize an *eslB* deletion strain, we found that the absence of EslB not only affects lysozyme resistance, but also endogenous cell lysis, cell wall biosynthesis, cell division and the ability of the bacterium to grow in media containing high concentrations of sugars. Our results indicate that EslB is by a yet unknown mechanism an important determinant for cell wall integrity in *L. monocytogenes*.

## INTRODUCTION

Gram-positive bacteria are surrounded by a complex cell wall, which is composed of a thick layer of peptidoglycan (PG) and cell wall polymers. The bacterial cell wall is important for the maintenance of cell shape, the ability of bacteria to withstand harsh environmental conditions and to prevent cell lysis (1, 2). Due to its importance, cell wall-targeting antibiotics such as β-lactam, glycopeptide and fosfomycin antibiotics are commonly used to treat bacterial infections (3, 4). These cell-wall targeting antibiotics inhibit enzymes involved in different stages of the PG biosynthesis process or sequester substrates of these enzymes (4). Moenomycin, another cell wall-targeting antibiotic, and β-lactam antibiotics, for instance, block the glycosyltransferase and transpeptidase activity of penicillin binding proteins, respectively, which are required for the polymerization and crosslinking of the glycan strands (5–7). Peptidoglycan is also the target of the cell wall hydrolase lysozyme, which is a component of animal and human secretions such as tears and mucus. Lysozyme cleaves the glycan strands of PG by hydrolysing the 1,4-β-linkage between *N*-acetylmuramic acid (MurNAc) and *N*-acetylglucosamine (GlcNAc). This reaction leads to a loss of cell integrity and results in cell lysis (8). The intracellular human pathogen *Listeria monocytogenes* is intrinsically resistant to lysozyme due to modifications of its PG. The *N*-deacetylase PgdA deacetylates GlcNAc residues, whereas MurNAc residues are acetylated by the *O*-acetyltransferase OatA (9, 10). Consequently, deletion of either of these enzymes results in reduced lysozyme resistance (9, 10). One or both of these enzymes are also present in other bacterial pathogens and important for lysozyme resistance, such as PgdA in *Streptococcus pneumoniae*, OatA in *Staphylococcus aureus* and PgdA and OatA in *Enterococcus faecalis* (11–14). Besides enzymes that directly alter the peptidoglycan structure, a number of other factors have been shown to contribute to lysozyme resistance in diverse bacteria. For instance, the cell wall polymer wall teichoic acid and the two-component system GraRS contribute to lysozyme resistance in *S. aureus* (15, 16). In *E. faecalis*, the extracytoplasmic function sigma factor SigV is required for the upregulation of *pgdA* expression in the presence of lysozyme (11, 17). Recently, some additional factors have been identified, which contribute to the intrinsic lysozyme resistance of *L. monocytogenes* such as the predicted carboxypeptidase PbpX, the transcription factor DegU and the noncoding RNA Rli31 (18). DegU and Rli31 are involved in the regulation of *pgdA* and *pbpX* expression in *L. monocytogenes* (18). Furthermore, components of a predicted ABC transporter encoded by the *lmo2769-6* operon in *L. monocytogenes* and here referred to as *eslABCR* for <u>e</u>longation, <u>s</u>ugar& and <u>l</u>ysozyme sensitive phenotype (Fig. 1) have been associated with lysozyme resistance (18–20). An *eslB* transposon insertion mutant was also shown to be more sensitive to cefuroxime and cationic antimicrobial peptides (18).

**Figure 1:**
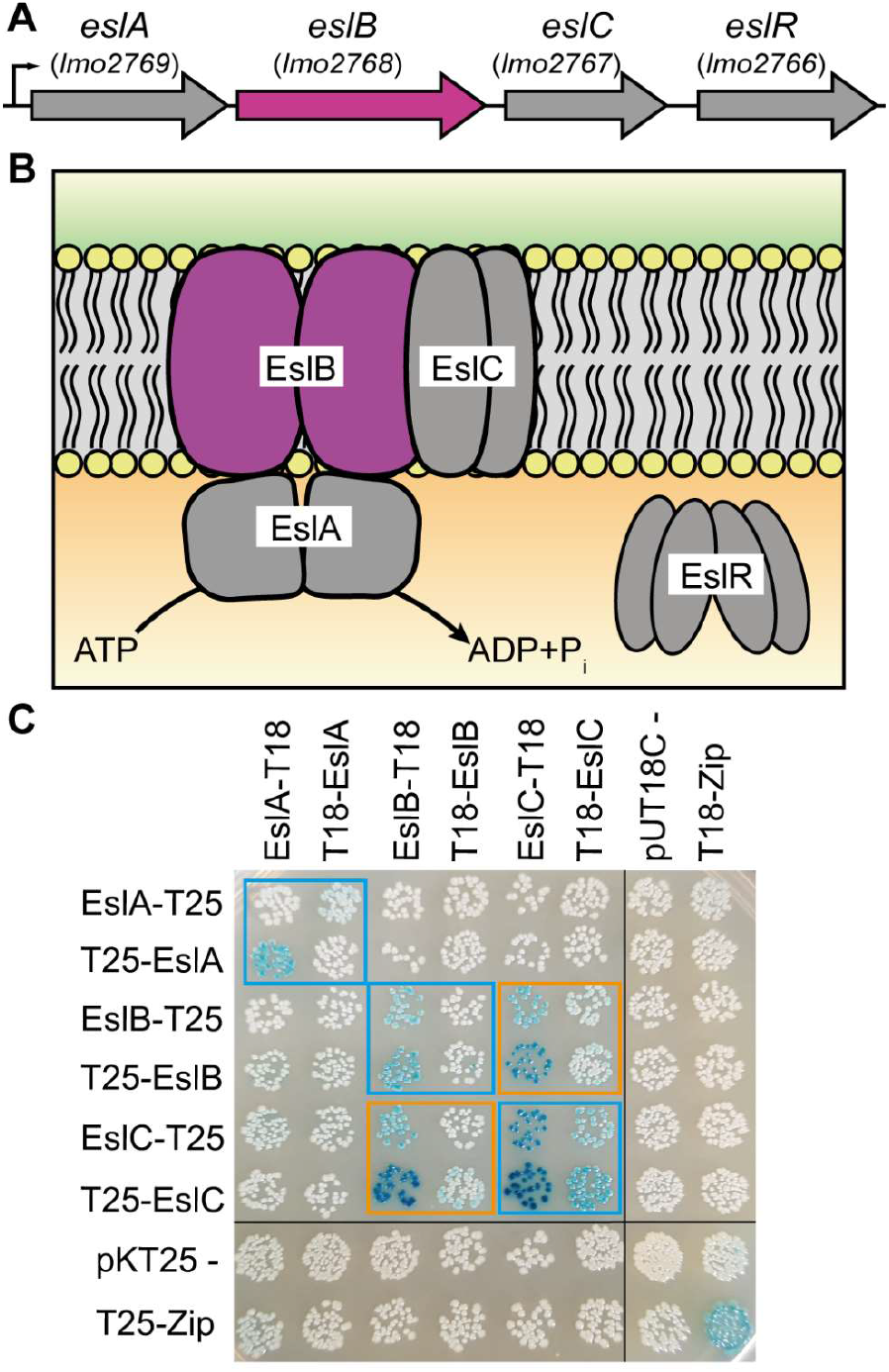
Schematic representation of the *L. monocytogenes eslABCR* operon and interaction of the ABC transporter components EslABC. (A) Genomic arrangement of the *eslABCR* operon in *L. monocytogenes*. Arrowheads indicate the orientation of the genes. Small black arrow indicates the promoter identified in a previous study (20). (B) Model of the ABC transporter composed of the NBD protein EslA, which hydrolyses ATP, the TMD proteins EslB and EslC, and the cytoplasmic RpiR family transcription regulator EslR. The *eslB* gene and EslB protein, which were investigated as part of the study, are highlighted in pink. (C) Interactions between the ABC transporter components. Plasmids encoding fusions of EslA, EslB and EslC and the T18- and T25-fragments of the *Bordetella pertussis* adenylate cyclase were co-transformed into *E. coli* BTH101. Empty vectors pKT25 and pUT18C were used as negative control and pKT25- and pUT18C-Zip as positive control. Black lines indicate where lanes, which were not required, were removed. Self-interactions are marked with blue boxes and protein-protein interactions with orange boxes. A representative image of three repeats is shown.

ABC transporters can either act as importers or exporters. Importers are involved in the uptake of sugars, peptides or other metabolites, which are recognized by substrate binding proteins. On the other hand, toxic compounds such as antibiotics can be exported by ABC exporters (21–23). They are usually composed of homo- or heterodimeric cytoplasmic nucleotide-binding domain (NBD) proteins, also referred to as ATP-binding cassette proteins, and homo- or heterodimeric transmembrane domain (TMD) proteins (24). In addition to NBDs and TMDs, ABC importers have an extracellular substrate binding protein (SBP) or a membrane-integrated S-component, which are important for the delivery of specific substrate molecules to the transporter or substrate binding, respectively (25–27). The *esl* operon encodes EslA, the NBD protein, EslB, the TMD protein forming part of the ABC transporter, EslC, a membrane protein of unknown function and EslR, a RpiR-type transcriptional regulator (Fig. 1). So far, it has not been investigated whether EslC is a component of the ABC transporter encoded in the *esl* operon. EslB and EslC could for instance interact with each other and form the transmembrane domain of the ABC transporter, or EslC could function independent from EslAB. Furthermore, it is not known whether the predicted ABC transporter EslABC acts as an importer or exporter and its exact cellular function has not been identified. Here, we show that the absence of EslB, one of the transmembrane components of the ABC transporter, leads to an increased lysozyme sensitivity due to an altered PG structure. In addition, deletion of *eslB* resulted in the production of a thinner cell wall, and thus to an increased endogenous cell lysis. Furthermore, cell division is perturbed in the absence of EslB. We hypothesize that EslB may be required for processes, which are important for the maintenance of the cell wall integrity of *L. monocytogenes* during stress conditions.

## MATERIALS AND METHODS

### Bacterial strains and growth conditions

All strains and plasmids used in this study are listed in Table S1. *Escherichia coli* strains were grown in Luria-Bertani (LB) medium and *Listeria monocytogenes* strains in brain heart infusion (BHI) medium at 37°C unless otherwise stated. If necessary, antibiotics and supplements were added to the medium at the following concentrations: for *E. coli* cultures, ampicillin (Amp) at 100 μg/ml, chloramphenicol (Cam) at 20 μg/ml and kanamycin (Kan) at 30 μg/ml, and for *L. monocytogenes* cultures, Cam at 10 μg/ml, erythromycin (Erm) at 5 μg/ml, Kan at 30 μg/ml, nalidixic acid (Nal) at 20 μg/ml, streptomycin (Strep) at 200 μg/ml and IPTG at 1 mM.

### Strain and plasmid construction

All primers used in this study are listed in Table S2. For the markerless in-frame deletion of *lmo2768* (*lmrg_01927*, *eslB*), approximately 1kb-DNA fragments up- and downstream of the *eslB* gene were amplified by PCR using primer pairs ANG2532/2533 and ANG2534/2535. The resulting PCR products were fused in a second PCR using primers ANG2532/2535, the product cut with BamHI and XbaI and ligated with plasmid pKSV7 that had been cut with the same enzymes. The resulting plasmid pKSV7-Δ*eslB* was recovered in *E. coli* XL1-Blue yielding strain ANG4236. The plasmid was subsequently transformed into *L. monocytogenes* strain 10403S and *eslB* deleted by allelic exchange using a previously described procedure (28). The deletion of *eslB* was verified by PCR. The deletion procedure was performed with two independent transformants and resulted in the construction of two independent *eslB* mutant strains 10403SΔ*eslB*_*(1)*_ (ANG4275) and 10403SΔ*eslB*_*(2)*_ (ANG5662). For complementation analysis, pIMK3-*eslB* was constructed, in which the expression of *eslB* can be induced by IPTG. The *eslB* gene was amplified using primer pair ANG2812/ANG2813, the product cut with NcoI and SalI and fused with pIMK3 that had been cut with the same enzymes. The resulting plasmid pIMK3-*eslB* was recovered in *E. coli* XL1-Blue yielding strain ANG4647. Due to difficulties in preparing electrocompetent cells of *L. monocytogenes eslB* mutant strains, plasmid pIMK3-*eslB* was first electroporated into the wildtype *L. monocytogenes* strain 10403S yielding strain 10403S pIMK3-*eslB* (ANG4678). In the second step, *eslB* was deleted from the genome of strain ANG4678 resulting in the construction of the first *eslB* complementation strain 10403SΔ*eslB*_*(1)*_ pIMK3-*eslB* (ANG4688, short 10403SΔ*eslB(1)* compl.). In addition, complementation plasmid pPL3e-P_*eslA*_-*eslABC* was constructed. To this end, the *eslABC* genes including the upstream promoter region were amplified by PCR using primers ANG3349/ANG3350. The resulting PCR product was cut with SalI and BamHI and fused with plasmid pPL3e that had been cut with the same enzymes. Plasmid pPL3e-P_*eslA*_-*eslABC* was recovered in *E. coli* XL1-Blue yielding strain ANG5660. Next, plasmid pPL3e-P_*eslA*_-*eslABC* was transformed into *E. coli* SM10 yielding strain ANG5661. Lastly, SM10 pPL3e-P_*eslA*_-*eslABC* was used as a donor strain to transfer plasmid pPL3e-P_*eslA*_-*eslABC* by conjugation into *L. monocytogenes* strain 10403SΔ*eslB*_*(2)*_ (ANG5662) using a previously described method (29). This resulted in the construction of the second *eslB* complementation strain 10403SΔ*eslB*_*(2)*_ pPL3e-P_*eslA*_-*eslABC* (ANG5663, short 10403SΔ*eslB*_*(2)*_ compl.). For the markerless in-frame deletion of *lmo2769* (*lmrg_01926, eslA*), and *lmo2767* (*lmrg_01928, eslC*), approximately 1kb-DNA fragments up- and downstream of the corresponding gene were amplified by PCR using primer pairs LMS160/161 and LMS159/162 (*eslA*), and LMS155/158 and LMS156/157 (*eslC*). The resulting PCR products were fused in a second PCR using primers LMS159/160 (*eslA*) and LMS155/156 (*eslC*). The products were cut with BamHI and EcoRI (*eslA*) and BamHI and KpnI (*eslC*) and ligated with plasmid pKSV7 that had been cut with the same enzymes. The resulting plasmids pKSV7-Δ*eslA* and pKSV7-Δ*eslC* were recovered in *E. coli* XL1-Blue yielding strains EJR54 and EJR43, respectively. The plasmids were subsequently transformed into *L. monocytogenes* strain 10403S and *eslA* and *eslC* deleted by allelic exchange yielding strains 10403SΔ*eslA* (LJR33) and 10403SΔ*eslC* (LJR7). Plasmid pPL3e-P_*eslA*_-*eslABC* was transferred into LJR33 and LJR7 via conjugation using strain SM10 pPL3e-P_*eslA*_-*eslABC* (ANG5661) as a donor strain, yielding strains 10403SΔ*eslA* pPL3e-P_*eslA*_-*eslABC* (LJR34, short 10403SΔ*eslA* compl.) and 10403SΔ*eslC* pPL3e-P_*eslA*_-*eslABC* (LJR21, short 10403SΔ*eslC* compl.).

For the construction of bacterial two hybrid plasmids, *eslA*, *eslB* and *eslC* were amplified by PCR using primer pairs JR44/45, JR46/47 and JR48/49, respectively. The resulting *eslA* and *eslC* fragments were cut with XbaI and KpnI and ligated into pKT25, pKNT25, pUT18 and pUT18C that had been cut with the same enzymes. The *eslB* fragment was cut with XbaI and BamHI and ligated into XbaI/BamHI cut pKT25, pKNT25, pUT18 and pUT18C. The resulting plasmids were recovered in *E. coli* XL1-Blue yielding strains XL1-Blue pKNT25-*eslA* (EJR4), XL1-Blue pKT25-*eslA* (EJR5), XL1-Blue pUT18-*eslA* (EJR6), XL1-Blue pUT18C-*eslA* (EJR7), XL1-Blue pKNT25-*eslB* (EJR8), XL1-Blue pKT25-*eslB* (EJR9), XL1-Blue pUT18-*eslB* (EJR10), XL1-Blue pUT18C-*eslB* (EJR11), XL1-Blue pKNT25-*eslC* (EJR12), XL1-Blue pKT25-*eslC* (EJR13) and XL1-Blue pUT18C-*eslC* (EJR15). Using this approach, we were unable to construct pUT18-*eslC* without acquiring mutations in *eslC*. In a second attempt to generate pUT18-*eslC*, plasmid pKT25-*eslC* (from strain EJR13) was cut with XbaI and KpnI, the *eslC* fragment extracted and ligated into XbaI/KpnI cut pUT18. The resulting plasmid was recovered in *E. coli* CLG190 yielding strain CLG190 pUT18-*eslC* (EJR14).

For the localization of an early cell division protein, the N-terminus of ZapA was fused to mNeonGreen. For this purpose, *mNeonGreen* and *zapA* genes were amplified using primer pairs JR73/JR39 and JR40/JR74, respectively. The resulting PCR products were fused in a second PCR using primers JR73/JR74, the product was cut with NcoI and SalI and ligated with pIMK2 that had been cut with the same enzymes. pIMK2-*mNeonGreen*-*zapA* was recovered in *E. coli* XL1-Blue and transformed into *E. coli* S17-1 yielding strains EJR39 and EJR60, respectively. S17-1 pIMK2-*mNeonGreen*-*zapA* was used as a donor strain to transfer the plasmid pIMK2-*mNeonGreen*-*zapA* by conjugation into *L. monocytogenes* strains 10403S (ANG1263) and 10403SΔ*eslB*_*(2)*_ (ANG5662) resulting in the construction of strains 10403S pIMK2-*mNeonGreen*-*zapA* (LJR28) and 10403SΔ*eslB*_*(2)*_ pIMK2-*mNeonGreen*-*zapA* (LJR29).

### Bacterial two-hybrid assays

Interactions between EslA, EslB and EslC were analyzed using bacterial adenylate cyclase two-hybrid (BACTH) assays (30). For this purpose, 15 ng of the indicated pKT25/pKNT25 and pUT18/pUT18C derivatives were co-transformed into *E. coli* strain BTH101. Transformants were spotted on LB agar plates containing 25 μg/ml kanamycin, 100 μg/ml ampicillin, 0.5 mM IPTG and 80 μg/ml X-Gal and the plates incubated at 30°C. Images were taken after an incubation of 48 h.

### Whole genome sequencing

Genomic DNA of *L. monocytogenes* was extracted using the FastDNA™ Kit (MP Biomedicals) and libraries for sequencing were prepared using the Illumina Nextera DNA kit. The samples were sequenced at the London Institute of Medical Sciences using an Illumina MiSeq instrument and a 150 paired end Illumina kit. The reads were trimmed, mapped to the *L. monocytogenes* 10403S reference genome (NC_017544) and single nucleotide polymorphisms (SNPs) with a frequency of at least 80% and small deletions (zero coverage) identified using the CLC workbench genomics (Qiagen).

### Growth analysis

*L. monocytogenes* strains were grown overnight in 5 ml BHI medium at 37°C with shaking. The next day, these cultures were used to inoculate 15 ml fresh BHI medium or BHI medium containing 0.5 M sucrose, fructose, glucose, maltose, galactose or sodium chloride to an OD_600_ of 0.05. The cultures were incubated at 37°C with shaking at 180 rpm, OD_600_ readings were taken every hour for 8 h.

### Determination of minimal inhibitory concentration (MIC)

The minimal inhibitory concentration for the cell wall-acting antibiotics penicillin and moenomycin and the cell wall hydrolase lysozyme was determined in 96-well plates using a microbroth dilution assay. Approximately 10^4^ *L. monocytogenes* cells were used to inoculate 200 μl BHI containing two-fold dilutions of the different antimicrobials. The starting antibiotic concentrations were: 0.025 μg/ml for penicillin G, 0.2 μg/ml for moenomycin and 10 mg/ml or 0.25 mg/ml for lysozyme. The 96-well plates were incubated at 37°C with shaking at 500 rpm in a plate incubator (Thermostar, BMG Labtech) and OD_600_ determined after 24 hours of incubation. The MIC value refers to the antibiotic concentration at which bacterial growth was inhibited by >90%.

### Plate spotting assay

Overnight cultures of the indicated *L. monocytogenes* strains were adjusted to an OD_600_ of 1 and serially diluted to 10^−6^. 5 μl of each dilution were spotted on BHI agar plates or BHI agar plates containing 100 μg/ml lysozyme, both containing 1 mM IPTG. Images of the plates were taken after incubating them for 20-24 h at 37°C.

### Peptidoglycan isolation and analysis

Overnight cultures of 10403SΔ*eslB*_*(1)*_ and 10403SΔ*eslB*_*(1)*_ compl. were diluted in 1 L BHI broth (supplemented with 1 mM IPTG for strain 10403SΔ*eslB*_*(1)*_ compl.) to an OD_600_ of 0.06 and incubated at 37°C. At an OD_600_ of 1, bacterial cultures were cooled on ice for 1h and the bacteria subsequently collected by centrifugation. The peptidoglycan was purified, digested with mutanolysin and the muropeptides analyzed by HPLC using an Agilent 1260 infinity system, as previously described (31, 32). Peptidoglycan of the wildtype *L. monocytogenes* strain 10403S was purified and analyzed in parallel. The chromatogram of the same wild-type control strain was recently published (33) and also used as part of this study, since all strains were analyzed at the same time. The major peaks 1-6 were assigned according to previously published HPLC spectra (18, 34), with peaks 2, 4, 5 and 6 corresponding to *N*-deacetylated GlcNAc residues. Peaks 1-2 correspond to monomeric and peaks 4-6 to dimeric (crosslinked) muropeptide fragments. The Agilent Technology ChemStation software was used to integrate the areas of the main muropeptide. For quantification, the sum of the peak areas was set to 100% and the area of individual peaks was determined. The sum of values for peaks 3-6 corresponds to the % crosslinking, whereas the deacetylation state was calculated by adding up the values for peaks 4, 5 and 6. Averages values and standard deviations were calculated from three independent extractions.

### *O*-acetylation assay

Peptidoglycan of strains 10403S, 10403SΔ*eslB*_*(1)*_ and 10403SΔ*eslB*_*(1)*_ compl., which had not been treated with hydrofluoric acid and alkaline phosphatase to avoid removal of the *O*-acetyl groups, was used for the *O*-acetylation assays. *O*-acetylation was measured colorimetrically according to the Hestrin method described previously (35) with slight modifications. Briefly, 800 μg of PG (dissolved in 500 μl H_2_O) were incubated with an equal volume of 0.035 M hydroxylamine chloride in 0.75 M NaOH for 10 min at 25°C. Next, 500 μl of 0.6 M of perchloric acid and 500 μl of 70 mM ferric perchlorate in 0.5 M perchloric acid were added. The color change resulting from the presence of *O*-acetyl groups was quantified at 500 nm. An assay reaction with 500 μl H_2_O was used as a blank for the absorbance measurement.

### Autolysis assays

*L. monocytogenes* strains were diluted in BHI or BHI medium supplemented with 0.5 M sucrose to an OD_600_ of 0.05 and grown for 4 h at 37°C. Cells were collected by centrifugation and resuspended in 50 mM Tris-HCl, pH 8 to an OD_600_ of 0.7-0.9 and incubated at 37°C. For penicillin- and lysozyme-induced lysis, 25 μg/ml penicillin G or 2.5 μg/ml lysozyme was added to the cultures. Autolysis was followed by determining OD_600_ readings every 15 min.

### Fluorescence and phase contrast microscopy

Overnight cultures of the indicated *L. monocytogenes* strains were diluted 1:100 in BHI medium and grown for 3 h at 37°C. For staining of the bacterial membrane, 100 μl of these cultures were mixed with 5 μl of 100 μg/ml nile red solution and incubated for 20 min at 37°C. The cells were washed twice with PBS and subsequently suspended in 50 μl of PBS. 1-1.5 μl of the different samples were subsequently spotted on microscope slides covered with a thin agarose film (1.5 % agarose in distilled water), air-dried and covered with a cover slip. Phase contrast and fluorescence images were taken at 1000x magnification using the Zeiss Axio Imager.A1 microscope coupled to an AxioCam MRm and processed using the Zen 2012 software (blue edition). The nile red fluorescence signal was detected using the Zeiss filter set 00. The length of 300 cells was measured for each experiment and the median cell length was calculated.

For ZapA-localization studies, overnight cultures of the indicated *L. monocytogenes* strains were grown in BHI medium at 37°C to an OD_600_ of 0.3-0.5. The staining of the bacterial membrane with nile red was performed as described above. After nile red staining, cells were fixed in 1.2% paraformaldehyde for 20 min at RT. 1-1.5 μl of the different samples were spotted on microscope slides as described above. Phase contrast and fluorescence images were taken at 1000x magnification using the Zeiss Axioskop 40 coupled to an AxioCam MRm and processed using the Axio Vision software (release 4.7). The nile red and mNeonGreen fluorescence signals were detected using the Zeiss filter set 43 and 37, respectively.

### Transmission electron microscopy

Overnight cultures of *L. monocytogenes* strains 10403S, 10403SΔ*eslB*_*(2)*_ and 10403SΔ*eslB*_*(2)*_ compl. were used to inoculate 25 ml BHI broth or BHI broth supplemented with 0.5 M sucrose to an OD_600_ of 0.05. Bacteria were grown at 37°C and 200 rpm for 3.5 h (BHI broth) or 6 h (BHI broth containing 0.5 M sucrose). 15 ml of the cultures were centrifuged for 10 min at 4000 rpm, the cell pellet washed twice in phosphate-buffered saline (127 mM NaCl, 2.7 mM KCl, 10 mM Na_2_HPO_4_, 1.8 mM KH_2_PO_4_, pH 7.4) and fixed overnight in 2.5 % (w/v) glutaraldehyde at 4°C. Cells were then mixed with 1.5 % (w/v, final concentration in PBS) molten Bacto-Agar, kept liquid at 55°C. After solidification, the agar block was cut into pieces with a volume of 1 mm^3^. A dehydration series was performed (15 % aqueous ethanol solution for 15 min, 30 %, 50 %, 70 % and 95 % for 30 min and 100 % for 2x 30 min) at 0°C, followed by an incubation step in 66 % (v/v, in ethanol) LR-white resin mixture (Plano) for 2 h at RT and embedded in 100 % LR-white solution overnight at 4°C. One agar piece was transferred to a gelatin capsule filled with fresh LR-white resin, which was subsequently polymerized at 55°C for 24 h. A milling tool (TM 60, Reichert & Jung, Vienna, Austria) was used to shape the gelatin capsule into a truncated pyramid. An ultramicrotome (Reichert Ultralcut E, Leica Microsystems, Wetzlar, Germany) and a diamond knife (Delaware Diamond Knives, Wilmington, DE, USA) were used to obtain ultrathin sections (80 nm) of the samples. The resulting sections were mounted on mesh specimen grids (Plano) and stained with 4 % (w/v) uranyl acetate solution (pH 7.0) for 10 min. Microscopy was performed using a Jeol JEM 1011 transmission electron microscope (Jeol Germany GmbH, Munich) at 80 kV. Images were taken at a magnification of 30,000 and recorded with an Orius SC1000 CCD camera (Pleasanton, CA, USA). For each replicate, 20 cells were photographed and cell wall thickness was measured at three different locations using the ImageJ software (36). The average of the three measurements was calculated and the average and standard deviation of 20 cells plotted. The experiment was performed twice.

### Cell culture

Bone marrow-derived macrophages (BMMs) were extracted from female C57BL/6 mice as described previously (37). BMMs were a gift from Charlotte S. C. Michaux and Sophie Helaine. 5×10^5^ BMMs were seeded per well of a 24-well plate and grown overnight in 500 μl high glucose Dulbecco’s Modified Eagle Medium (DMEM) at 37°C and 5% CO_2_. *L. monocytogenes* strains were grown overnight without shaking in 2 ml BHI medium at 30°C. The next morning, bacteria were opsonized with 8% mouse serum (Sigma-Aldrich) at room temperature for 20 min and BMMs were infected for one hour at a multiplicity of infection (MOI) of 2. BMMs were washed with PBS and 1 ml DMEM containing 40 μg/ml gentamycin was added to kill extracellular bacteria. After 1 h, cells were washed with PBS and covered with 1 ml DMEM containing 10 μg/ml gentamycin. The number of recovered bacteria was determined 2, 4, 6 and 8 h post infection. To this end, BMMs were lysed using 1 ml PBS containing 0.1% (v/v) triton X-100 and serial dilutions were plated on BHI agar plates. The number of colony forming units (CFUs) was determined after incubating the plates overnight at 37°C. Three technical repeats were performed for each experiment and average values calculated. Average values and standard deviations from three independent experiments were plotted.

### *Drosophila melanogaster* infections

Fly injections were carried out with microinjection needles produced from borosilicate glass capillaries (World Precision Instruments TW100-4) and a needle puller (Model PC-10, Narishige). Injections were performed using a Picospritzer III system (Parker Hannifin), and the injection volume was calibrated by expelling a drop of liquid from the needle into a pot of mineral oil and halocarbon oil (both Sigma). The expelled drop was measured using the microscope graticule to obtain a final injection volume of 50 nanolitres (nl). Flies were then anesthetized with CO_2_ and injected with either 50 nl of bacterial suspension in PBS or sterile PBS. 5-7-day old age matched male flies were used for all experiments. Flies were grouped into uninjected control, wounding control (injection with sterile PBS), and flies infected with *L. monocytogenes*. Each group consisted of 58-60 flies. All survival experiments were conducted at 29°C. Dead flies were counted daily. Food vials were placed horizontally to reduce the possibility of fly death from flies getting stuck to the food, and flies were transferred to fresh food every 3-4 days. For the quantification of the bacterial load, 16 flies per condition and per bacterial strain were collected at the indicated time points. The flies were homogenised in 100 μl of TE-buffer pH 8 containing 1% Triton X-100 and 1% Proteinase K (NEB, P8107S). Homogenates were incubated for 3 h at 55°C followed by a 10 min incubation step at 95°C. Following incubation, qPCR was carried out using the *actA* gene specific primers EGD-E_ActA_L1 and EGD-E_ActA_R1 to determine the number of bacterial colony forming units. PCR was performed with Sensimix SYBR Green no-ROX (Bioline) on a Corbett Rotor-Gene 6000. The cycling conditions were as follows: Hold 95°C for 10 min, then 45 cycles of 95°C for 15 s, 57°C for 30 s, 72°C for 30 s, followed by a melting curve. Gene abundances were calculated as previously described (38).

## RESULTS

### EslC interacts with the transmembrane protein EslB

Previously it has been shown that *L. monocytogenes* strains with mutations in the *eslABCR* operon (Fig. 1A) display decreased resistance towards the cell wall hydrolase lysozyme (18, 19). The *esl* operon encodes the ATP binding protein EslA and the transmembrane proteins EslB and EslC, which are proposed to form an ABC transporter. However, it is currently unknown if EslC forms part of the ABC transporter as depicted in Figure 1B and if it is required for the function of the transporter. To gain insights into the composition of the ABC transporter, we assessed the interaction between EslA, EslB and EslC using the bacterial adenylate cyclase-based two-hybrid system. In addition to self-interactions of EslA, EslB and EslC, we observed an interaction between EslB and EslC (Fig. 1C), indicating that EslC might be part of the ABC transporter.

### Deletion of *eslB* in *L. monocytogenes* leads to lysozyme sensitivity and an altered peptidoglycan structure

An *eslA* in-frame deletion mutant and an *eslB* transposon insertion mutant were shown to be more sensitive to lysozyme compared to the wildtype strain (18, 19). However, it is still unknown how the function of an ABC transporter is linked to this phenotype. To investigate this further, strains with markerless in-frame deletions in *eslA*, *eslB* and *eslC* were constructed in the *L. monocytogenes* strain background 10403S. First, the lysozyme resistance of these mutants was assessed using a plate spotting assay. The *eslA* and *eslB* mutants showed reduced growth on BHI plates containing 100 μg/ml lysozyme compared to the wildtype and *eslA* and *eslB* complementation strains (Fig. 2A). On the other hand, no phenotype was observed for the *eslC* mutant (Fig. 2A). Since deletion of *eslA* and *eslB* resulted in a decreased lysozyme resistance, and an *eslA* mutant has already been characterized in previous work (19), we focused here on the characterization of the *eslB* deletion strain.

**Figure 2:**
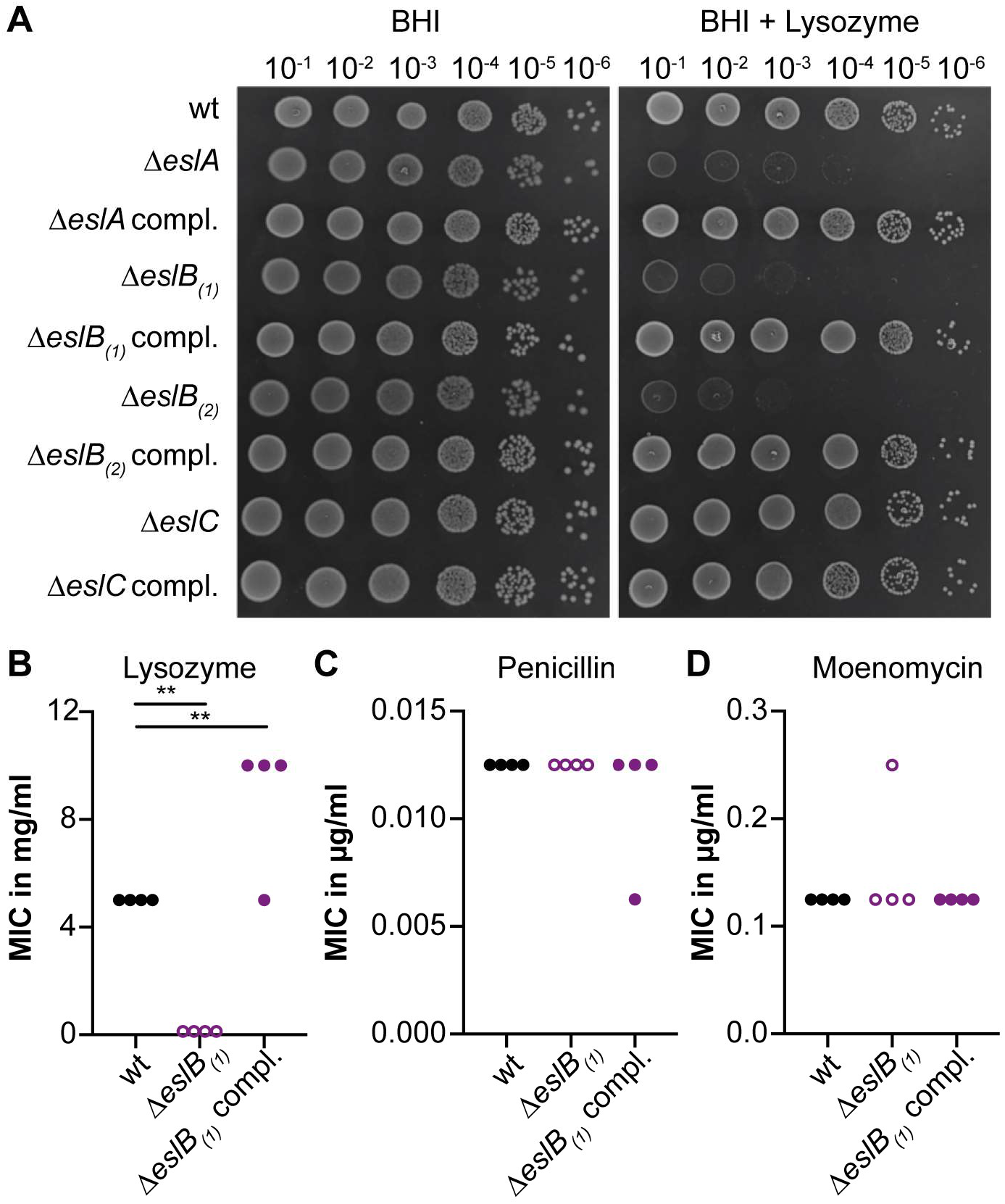
Impact of *eslB* deletion on resistance towards cell wall-targeting antimicrobials. (A) Plate spotting assay. Dilutions of overnight cultures of *L. monocytogenes* strains 10403S (wt), 10403SΔ*eslA*, 10403SΔ*eslA* compl., 10403SΔ*eslB*_*(1)*_, 10403SΔ*eslB*_*(1)*_ compl., 10403SΔ*eslB*_*(2)*_, 10403SΔ*eslB*_*(2)*_ compl., 10403SΔ*eslC*, and 10403SΔ*eslC* compl. were spotted on BHI plates and BHI plates containing 100 μg/ml lysozyme, both supplemented with 1 mM IPTG. A representative result from three independent experiments is shown. (B-D) Minimal inhibitory concentration (MIC) of *L. monocytogenes* strains 10403S (wt), 10403SΔ*eslB*_*(1)*_ and 10403SΔ*eslB*_*(1)*_ compl. towards (B) lysozyme, (C) penicillin G and (D) moenomycin. Strain 10403SΔ*eslB*_*(1)*_ compl. was grown in the presence of 1 mM IPTG. The results of four independent experiments are shown. For statistical analysis, a one-way ANOVA followed by a Dunnett’s multiple comparisons test was used (** *p*≤0.01).

In the course of the study, we determined the genome sequence of the originally constructed *eslB* mutant (10403SΔ*eslB*_*(1)*_) by whole genome sequencing (WGS) and identified an additional small deletion in gene *lmo2396* coding for an internalin protein with a leucine-rich repeat (LRR) and a mucin-binding domain (Table S3). While to the best of our knowledge, the contribution of Lmo2396 to the growth and pathogenicity of *L. monocytogenes* has not yet been investigated, other internalins are important and well-established virulence factors (39, 40). Our WGS analysis also revealed a single point mutation in gene *lmo2342*, coding for a pseudouridylate synthase in the complementation strain 10403SΔ*eslB*_*(1)*_ compl. (Table S3). Since we identified an additional mutation in a gene coding for a potential virulence factor in the *eslB* mutant, we constructed a second independent *eslB* mutant, 10403SΔ*eslB*_*(2)*_. We also constructed a second complementation strain, strain 10403SΔ*eslB*_*(2)*_ *PeslA-eslABC* (or short 10403SΔ*eslB*_*(2)*_ compl.), in which the *eslABC* genes are expressed from the native *eslA* promoter from a chromosomally integrated plasmid. Our WGS analysis revealed that strain 10403SΔ*eslB*_*(2)*_ did not contain any secondary mutations (Table S3). A 1-bp deletion in gene *lmo2022* encoding a predicted NifS-like protein required for NAD biosynthesis, was identified in strain 10403SΔ*eslB*_*(2)*_ compl. (Table S3), which if non-complementable phenotypes are observed needs to be kept in mind. We confirmed that our second *eslB* mutant strain 10403SΔ*eslB*_*(2)*_ showed the same lysozyme sensitivity phenotype and that this phenotype could be complemented in strain 10403SΔ*eslB*_*(2)*_ compl., in which *eslB* is expressed along with *eslA and eslC* from its native promoter (Fig. 2A). Since we only identified the genomic alterations in the course of the study, some experiments were performed as stated in the text with the original *eslB* mutant and complementation strains 10403SΔ*eslB*_*(1)*_ and 10403SΔ*eslB*_*(1)*_ compl., while other experiments were conducted with strains 10403SΔ*eslB*_*(2)*_ and 10403SΔ*eslB*_*(2)*_ compl.

Using microbroth dilution assays, we observed a 40-fold lower MIC for lysozyme for *L. monocytogenes* strain 10403SΔ*eslB*_*(1)*_ as compared to the wildtype strain (Fig. 2B and S1A) (18, 19). This phenotype could be complemented and strain 10403SΔ*eslB*_*(1)*_ compl., in which *eslB* is expressed from an IPTG-inducible promoter, is even slightly more resistant to lysozyme as compared to the wildtype strain (Fig. 2B). Next, we tested whether the resistance towards two cell wall-targeting antibiotics, namely penicillin and moenomycin, is changed upon deletion of *eslB*. The MIC values obtained for the wildtype, *eslB* deletion and *eslB* complementation strains were comparable (Fig. 2C-D), suggesting that the deletion of *eslB* does not lead to a general sensitivity to all cell wall-acting antimicrobials but is specific to lysozyme. In *L. monocytogenes*, lysozyme resistance is achieved by the modification of the peptidoglycan (PG) by *N*-deacetylation via PgdA and *O*-acetylation via OatA (9, 10). To assess whether deletion of *eslB* affects the *N*-deacetylation and crosslinking of PG, PG was isolated from wildtype 10403S, the *eslB* deletion and complementation strains, digested with mutanolysin and the muropeptides analyzed by high performance liquid chromatography (HPLC). This analysis revealed a slight increase in PG crosslinking in the *eslB* mutant strain (68±0.53%) compared to the wildtype (65.47±0.31%) and the complementation strain grown in the presence of IPTG (64.57±2.3%) (Fig. 3A-B). The GlcNAc residues of the PG isolated from the *eslB* deletion strain were also slightly more deacetylated (71.54±0.21%) as compared to the wildtype (67.17±0.31%) and the complementation strain (67±2.27%) (Fig. 3A-B), which should theoretically result in an increase and not decrease in lysozyme resistance. However, when we assessed the degree of *O*-acetylation using a colorimetric assay, the PG isolated from the *eslB* mutant was less *O*-acetylated compared to the wildtype and the complementation strain (Fig. 3C). Taken together, our results suggest that slight changes in the PG structure and in particular the observed reduction in *O*-acetylation likely contribute to the lysozyme sensitivity of the *eslB* deletion strain.

**Figure 3:**
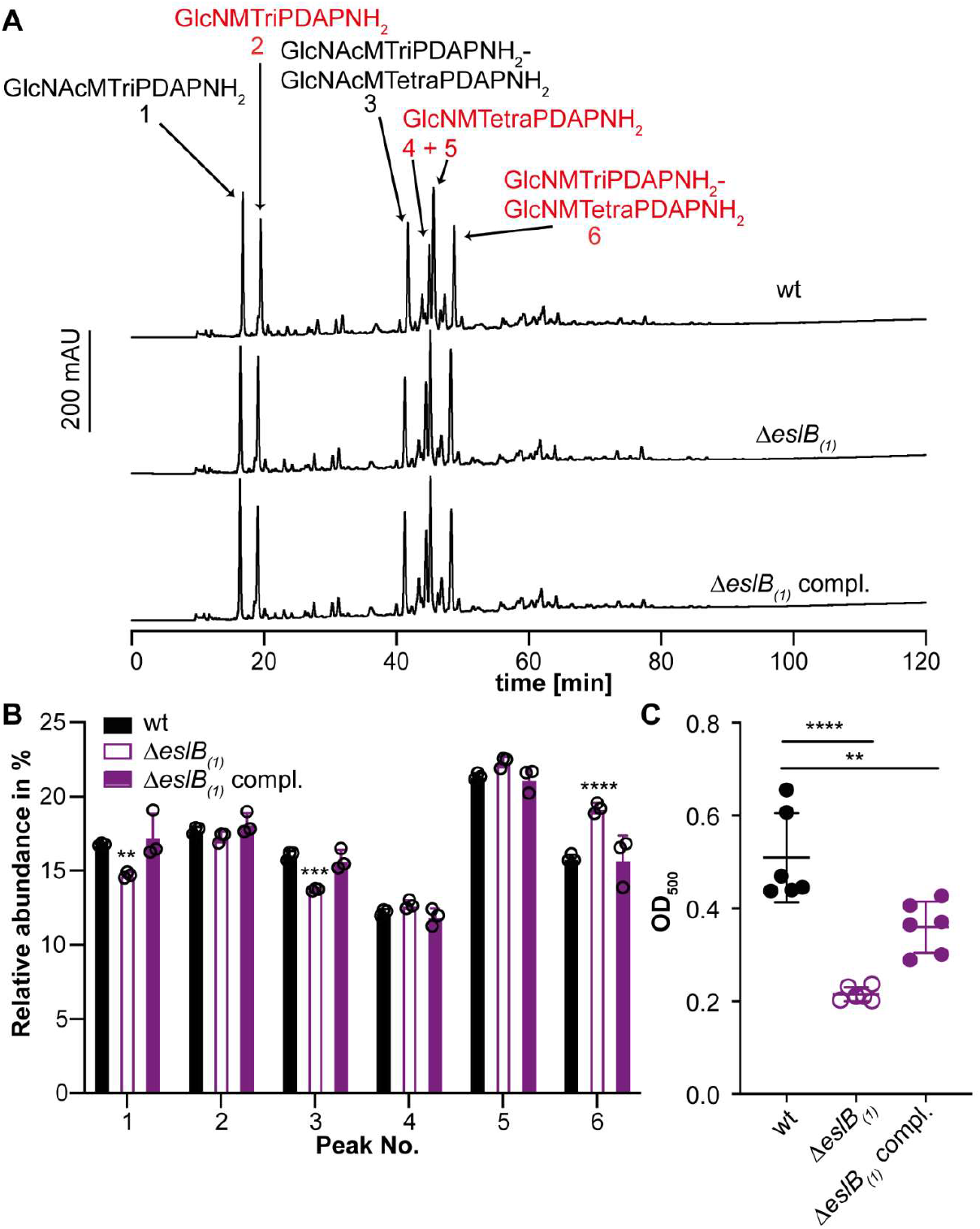
Deletion of *eslB* leads to changes in the peptidoglycan structure. (A) HPLC analysis of muropeptides derived from mutanolysin digested peptidoglycan isolated from strains 10403S (wt), 10403SΔ*eslB*_*(1)*_ and 10403SΔ*eslB*_*(1)*_ compl.. The muropeptide spectrum of the wildtype strain 10403S has been previously published (33). Major muropeptide peaks are labelled and numbered 1-6 according to previously published HPLC spectra (18, 34), with labels shown in red corresponding to muropeptides with *N*-deacetylated GlcNAc residues and peaks 1-2 corresponding to monomeric and 4-6 to dimeric (crosslinked) muropeptide fragments. Muropeptide abbreviations: GlcNAc – *N*-acetylglucosamine; GlcN – glucosamine; M – *N*-acetylmuramic acid; TriPDAPNH2 – L-alanyl-γ-D-glutamyl-amidated *meso*-diaminopimelic acid; TetraPDAPGlcNAc - L-alanyl-γ-D-glutamyl-amidated *meso*-diaminopimelyl-D-alanine. (B) Quantification of the relative abundance of muropeptide peaks 1-6 for peptidoglycan isolated of strains 10403S (wt), 10403SΔ*eslB*_*(1)*_ and 10403SΔ*eslB*_*(1)*_ compl.. For quantification, the sum of the peak areas was set to 100% and the area of individual peaks was determined. Average values and standard deviations were calculated from three independent peptidoglycan extractions and plotted. For statistical analysis, a two-way ANOVA followed by a Dunnett’s multiple comparisons test was used (** *p*≤0.01, *** *p*≤0.001, **** *p*≤0.0001). (C) The degree of *O*-acetylation of purified peptidoglycan of strains 10403S (wt), 10403SΔ*eslB*_*(1)*_ and 10403SΔ*eslB*_*(1)*_ compl. was determined by a colorimetric assay as described in the methods section. Average values and standard deviations were calculated from three independent peptidoglycan extractions and two technical repeats and plotted. For statistical analysis, a two-way ANOVA followed by a Dunnett’s multiple comparisons test was used (** *p*≤0.01, **** *p*≤0.0001).

### Deletion of *eslB* results in a growth defect in high sugar media

The bacterial cell wall is an important structure to maintain the cell integrity and to prevent lysis due to high internal turgor pressure or when bacteria experience changes in the external osmolality. Alterations of the PG structure or other cell wall defects leading to an impaired cell wall integrity could affect the growth of bacteria in environments with high osmolalities, e.g. in the presence of high salt or sugar concentrations. Next, we compared the growth of the wildtype, the *eslB* mutant and complementation strains at 37°C in different media. No growth difference could be observed between the strains tested, when grown in BHI medium (Fig. 4A and S1B). However, the *eslB* deletion strain grew slower in BHI medium containing 0.5 M sucrose as compared to the wildtype and the *eslB* complementation strain (Fig. 4B and S1C). A similar growth phenotype could be observed when the strains were grown in BHI medium containing either 0.5 M fructose, glucose, maltose or galactose (Fig. S2). In contrast, the presence of 0.5 M NaCl did not affect the growth of the *eslB* deletion strain (Fig. 4C). These results suggest that the observed growth defect seen for the *eslB* mutant is not solely caused by the increase in external osmolality, but rather seems to be specific to the presence of high concentrations of sugars.

**Figure 4:**
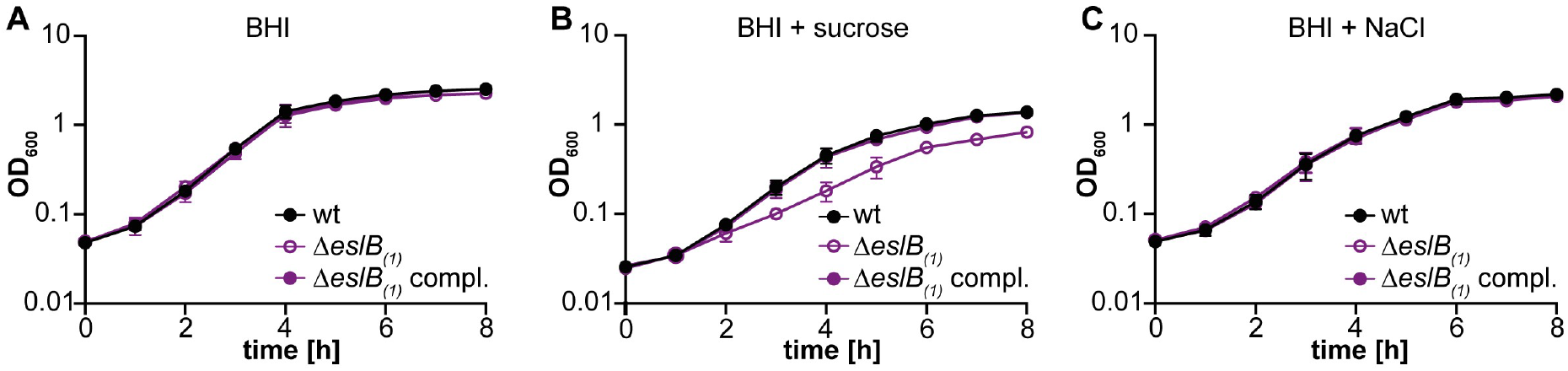
Addition of sucrose but not NaCl negatively impacts the growth of the *L. monocytogenes eslB* mutant strain. (A-C). Bacterial growth curves. *L. monocytogenes* strains 10403S (wt), 10403SΔ*eslB*_*(1)*_ and 10403SΔ*eslB*_*(1)*_ compl. were grown in (A) BHI broth, (B) BHI broth containing 0.5 M sucrose or (C) BHI broth containing 0.5 M NaCl. Strain 10403SΔ*eslB*_*(1)*_ compl. was grown in the presence of 1 mM IPTG. OD_600_ readings were determined at hourly intervals and the average values and standard deviations from three independent experiments calculated and plotted.

### Deletion of *eslB* results in increased endogenous and lysozyme-induced lysis

The observed lysozyme sensitivity and the growth defect of the *eslB* deletion strain in media containing high concentrations of sucrose raised the question, whether the absence of EslB might also cause an impaired cell wall integrity and an increased autolysis due to this impairment. To test this, autolysis assays were performed. To this end, the *L. monocytogenes* wildtype strain 10403S, the *eslB* deletion and complementation strains were grown in BHI medium and subsequently transferred in a Tris-HCl buffer (pH 8). After 2 h incubation at 37°C, the OD_600_ of the suspensions of the wildtype and *eslB* complementation strain had dropped to 89.9±1.6% and 86.5±2.9% of the initial OD_600_, respectively (Fig. 5A). Enhanced endogenous cell lysis was observed for the *eslB* mutant strain and the OD_600_ of the suspensions dropped to 68.8±1.7% within 2 h (Fig. 5A). The addition of penicillin had no impact on the cell lysis of any of the strains tested (Fig. 5B). On the other hand, the addition of 2.5 μg/ml lysozyme increased the rate of cell lysis of all strains, but had a particularly drastic effect on the *eslB* mutant. After 30 min, the OD_600_ reading of a suspension of the *eslB* deletion strain had dropped to 50.3±10.2%. For the wildtype and *eslB* complementation strains, it took 90 min to see a 50% reduction in the OD_600_ readings (Fig. 5C).

**Figure 5:**
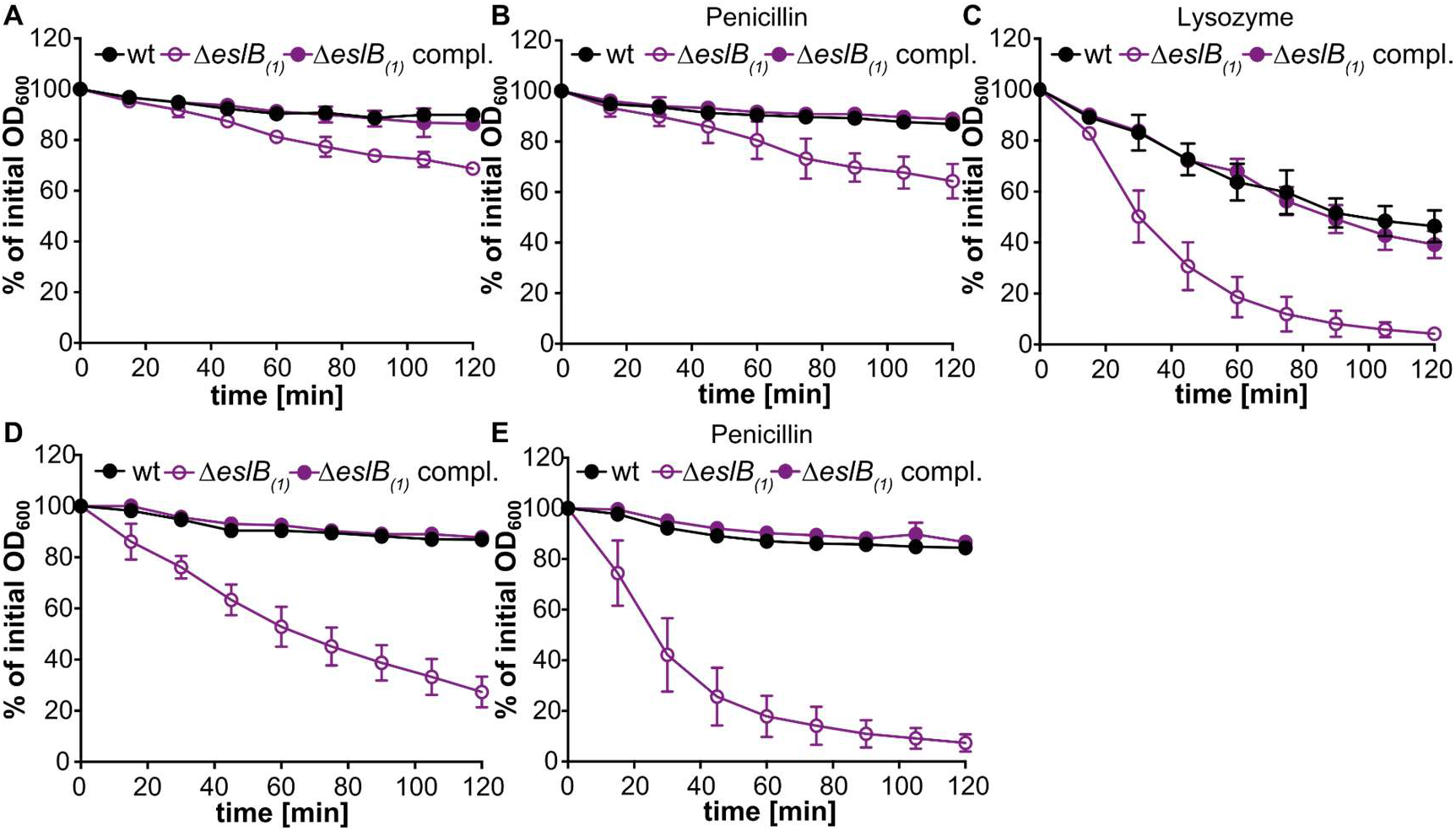
An *L. monocytogenes eslB* deletion strain shows increased endogenous and lysozyme-induced autolysis. Autolysis assays were performed with *L. monocytogenes* strains 10403S (wt), 10403SΔ*eslB*_*(1)*_ and 10403SΔ*eslB*_*(1)*_ compl.. Bacteria were grown for 4 h in (A-C) BHI medium or (D-E) in BHI medium containing 0.5 M sucrose (containing 1 mM IPTG for 10403SΔ*eslB*_*(1)*_ compl.) and subsequently bacterial suspensions prepared in (A, D) 50 mM Tris HCl pH 8, (B, E) 50 mM Tris HCl pH 8 containing 25 μg/ml penicillin, or (C) 2.5 μg/ml lysozyme. Cell lysis was followed by taking OD_600_ readings every 15 min. The initial OD_600_ reading for each bacterial suspension was set to 100% and subsequent readings are shown as % of the initial OD_600_ reading. The average % OD_600_ values and standard deviations were calculated from three independent experiments and plotted.

Next, we wanted to determine what impact the growth in the presence of high levels of sucrose has on endogenous bacterial autolysis rates. To this end, the wildtype 10403S, *eslB* mutant and complementation strains were grown in BHI medium supplemented with 0.5 M sucrose, cell suspensions prepared in Tris-buffer and used in autolysis assays. While the wildtype and complementation strain showed similar autolysis rates following growth in BHI sucrose medium (Fig. 5D) as after growth in BHI medium (Fig 5A), the *eslB* mutant lysed rapidly following growth in BHI 0.5 M sucrose medium (Fig. 5E). The lysis of the *eslB* mutant strain could be further enhanced by the addition of 25 μg/ml penicillin, a concentration which only acts bacteriostatic on the wildtype *L. monocytogenes* strain 10403S (Fig. 5E). These findings indicate that the *eslB* mutant is sensitive to osmotic downshifts and we thus wondered whether in addition to the changes in the PG modifications and crosslinking, more general differences in the ultrastructure of the cell wall might be observed. To test this, cells of *L. monocytogenes* strains 10403S, 10403SΔ*eslB*_*(2)*_ and 10403SΔ*eslB*_*(2)*_ compl. were subjected to transmission electron microscopy. The *eslB* deletion strain produces a thinner PG layer of 15.8±1.9 nm, when grown in BHI broth as compared to the wildtype (20±3.4 nm) and the complementation strain (20±4.3 nm, Fig. 6A-B). This phenotype was even more pronounced when the strains were grown in BHI broth containing 0.5 M sucrose. The PG layer of the *eslB* mutant had a thickness of 15±2 nm, while wildtype and the complementation strain produced a PG layer of 21.4±3.1 and 23.3±2.8 nm, respectively (Fig. 6A-B). We hypothesize that the enhanced endogenous lysis of the *eslB* mutant is likely caused by a thinner PG layer combined with the observed alterations in PG structure such as reduced *O-*acetylation.

**Figure 6:**
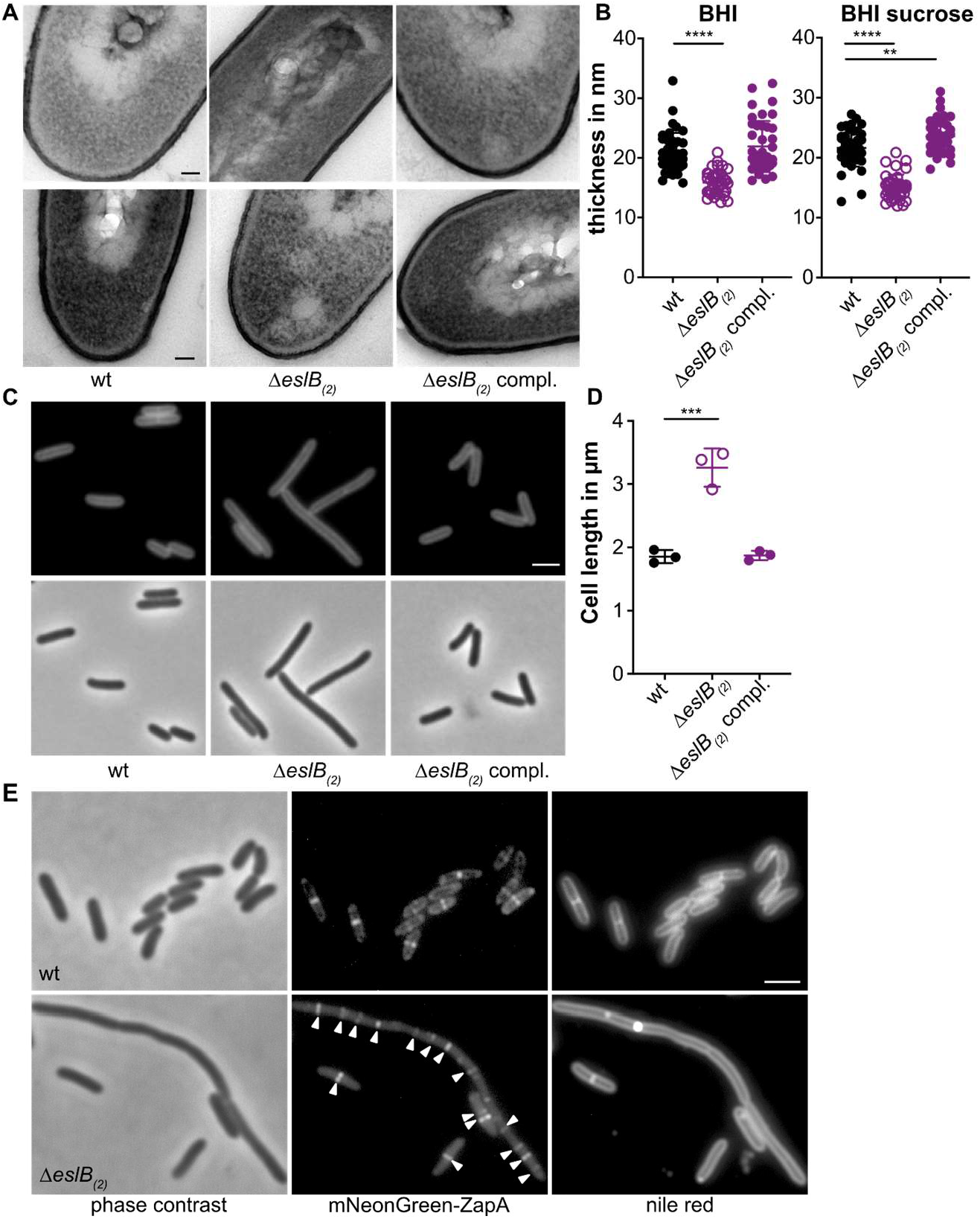
The *L. monocytogenes eslB* mutant produces a thinner cell wall, has a cell division defect and bacteria have an increased cell length. (A) Transmission electron microscopy images. Ultrathin-sectioned cells of *L. monocytogenes* strains 10403S (wt), 10403SΔ*eslB*_*(2)*_ and 10403SΔ*eslB*_*(2)*_ compl. were subjected to transmission electron microscopy after growth in BHI broth (upper panel) or BHI broth containing 0.5 M sucrose (lower panel). Scale bar is 50 nm. Representative images from two independent experiments are shown. (B) Cell wall thickness. Per growth condition, cell wall thickness of 40 cells was measured at three different locations and the average values plotted. For statistical analysis, a two-way ANOVA followed by a Dunnett’s multiple comparisons test was used (** *p*≤0.01, **** *p*≤0.0001). (C) Microscopy images of *L. monocytogenes* strains 10403S (wt), 10403SΔ*eslB*_*(2)*_ and 10403SΔ*eslB*_*(2)*_ compl.. Bacterial membranes were stained with nile red and cells analyzed by phase contrast and fluorescence microscopy. Scale bar is 2 μm. Representative images from three independent experiments are shown. (D) Cell length of *L. monocytogenes* strains 10403S (wt), 10403SΔ*eslB*_*(2)*_ and 10403SΔ*eslB*_*(2)*_ compl.. The cell length of 300 cells per strain was measured and the median cell length calculated. Three independent experiments were performed, and the average values and standard deviation of the median cell length plotted. For statistical analysis, a one-way ANOVA analysis followed by a Dunnett’s multiple comparisons test was used (*** *p*≤0.001). (E) Localization of mNeonGreen-ZapA in *L. monocytogenes* strains 10403S (wt) and 10403SΔ*eslB*_*(2)*_. Bacterial membranes were stained with nile red and cells analyzed by phase contrast (left panel) and fluorescence microscopy to detect mNeonGreen (middle panel) and nile red fluorescence signals (right panel). White arrows highlight ZapA foci in cells of the *L. monocytogenes eslB* mutant. Scale bar is 2 μm. Representative images from three independent experiments are shown.

### The *eslB* deletion strain is impaired in cell division, but not in virulence

The increased endogenous autolysis together with the observed changes in the PG structure of the *eslB* deletion strain could result in an increased sensitivity to autolysins. The major autolysins of *L. monocytogenes* are p60 and NamA, which hydrolyze PG and are required for daughter cell separation during cell division (41, 42). Absence of either p60 or NamA results in the formation of chains (41, 42). We thus wondered whether deletion of *eslB* causes changes in the cell morphology of *L. monocytogenes*. Microscopic analysis revealed that cells lacking EslB are significantly longer with a median cell length of 3.26±0.25 μm as compared to the *L. monocytogenes* wildtype strain, which produced cells with a length of 1.85±0.08 μm (Fig. 6C-D), highlighting that the absence of EslB results in a cell division defect. To test whether the assembly of the early divisome is affected by the absence of EslB, we compared the localization of the early cell division protein ZapA in the wildtype and the *eslB* mutant background. In *L. monocytogenes* wildtype cells, a signal was observed at midcell for cells, which have initiated the division process (Fig. 6E). While short cells of the *eslB* mutant also only possess a single fluorescent signal, several ZapA fluorescence foci could be observed in elongated cells (Fig. 6E), suggesting that early cell division proteins can still localize in the *eslB* mutant and that a process downstream seems to be perturbed in the absence of EslB.

Next, we wanted to assess whether the impaired cell integrity and the observed cell division defect would also affect the virulence of the *L. monocytogenes eslB* mutant. Of note, in a previous study, it was shown that deletion of *eslA*, coding for the ATP-binding protein component of the ABC transporter, has no effect on the cell-to-cell spread of *L. monocytogenes* (19). To determine whether EslB is involved in the virulence of *L. monocytogenes*, primary mouse macrophages were infected with wildtype 10403S, the *eslB* mutant 10403SΔ*eslB*_*(2)*_ and complementation strain 10403SΔ*eslB*_*(2)*_ compl.. All three strains showed a comparable intracellular growth pattern (Fig. 7A), suggesting that EslB does not impact the ability of *L. monocytogenes* to grow in primary mouse macrophages. Next, we assessed the ability of the *eslB* deletion strains to kill *Drosophila melanogaster* as lysozyme is one important component of its innate immune response (43). All uninfected flies (U/C) and 96.6% of the flies that were injected with PBS survived the duration of the experiment (Fig. 7B). No statistically significant difference could be observed for the survival and bacterial load of flies infected with the different *L. monocytogenes* strains (Fig. 7B-C). These results indicate that, while EslB does not impact the ability of *L. monocytogenes* to infect and kill mammalian macrophages or *Drosophila melanogaster*, it nonetheless impacts the cell division and cell wall integrity of *L. monocytogenes* and consistent with this we have identified changes in the composition and thickness of the peptidoglycan layer.

**Figure 7:**
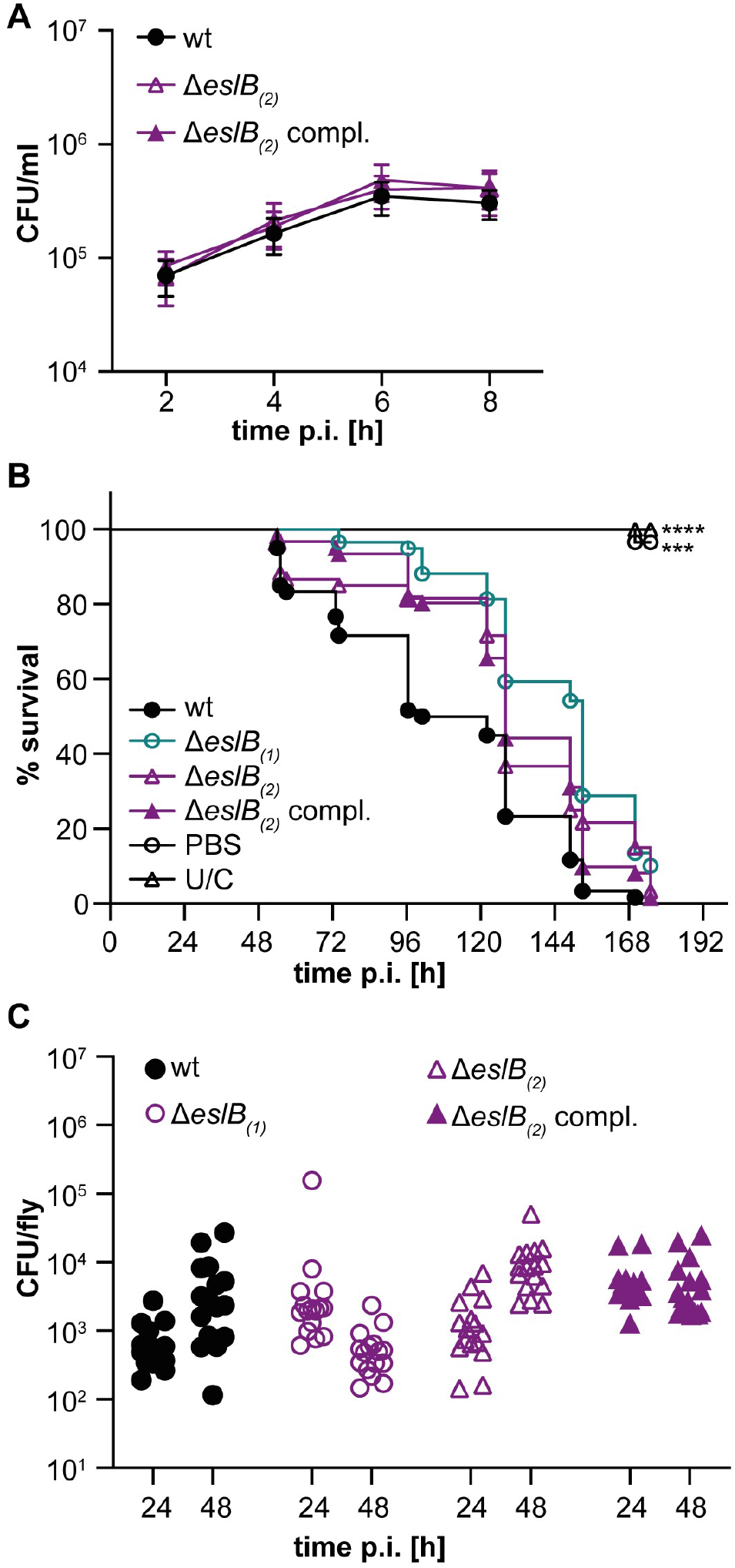
Impact of the deletion of *eslB* on the virulence of *L. monocytogenes*. (A) Intracellular growth of *L. monocytogenes* strains 10403S (wt), 10403SΔ*eslB*_*(2)*_ and 10403SΔ*eslB*_*(2)*_ compl. in mouse bone marrow-derived macrophages (BMMs). The infection assay was performed as described in the methods section. The average CFU count/ml and standard deviations from three independent experiments were calculated and plotted. (B) Survival curve of flies infected with *L. monocytogenes*. Flies were infected with *L. monocytogenes* strains 10403S (wt), 10403SΔ*eslB*_*(1)*_, 10403SΔ*eslB*_*(2)*_ and 10403SΔ*eslB*_*(2)*_ compl.. Uninjected control flies (U/C) and flies injected with PBS were used as controls. Fly death was monitored daily. For statistical analysis, a one-way ANOVA followed by a Dunnett’s multiple comparisons test was used (*** *p*≤0.001, **** *p*≤0.0001). (C) Bacterial quantification. 16 flies infected with the indicated *L. monocytogenes* strain were collected 24 and 48 h post infection and bacterial load (CFU) determined as described in the methods section. For statistical analysis, a nested one-way ANOVA followed by a Dunnett’s multiple comparisons test was used. The observed differences were not statistically significant.

## DISCUSSION

Over the past years, several determinants contributing to the intrinsic lysozyme resistance of *L. monocytogenes* have been described (9, 10, 18, 19). One of these is a predicted ABC transporter encoded as part of the *eslABCR* operon (18, 19). In this study, we aimed to further investigate the role of the ABC transporter EslABC in lysozyme resistance of *L. monocytogenes*. Using bacterial two hybrid assays, we could show that EslB and EslC interact with each other and hence it is tempting to speculate that the transmembrane component of the ABC transporter consists of a heterodimer of EslB and EslC. However, analysis of different deletion mutants revealed that only EslA and EslB are required for lysozyme resistance of *L. monocytogenes*, suggesting that EslC is not required for the function of the ABC transporter under our assay conditions. Surprisingly, we did not observe an interaction between EslA and EslB using bacterial two hybrid assays, thus, further experiments are required to determine the composition of the ABC transporter and its interaction partners.

Next, we analyzed the PG structure of the *eslB* deletion strain and found that the PG isolated from the *eslB* mutant was slightly more crosslinked and also the fraction of deacetylated GlcNAc residues was slightly increased as compared to the PG isolated from the wildtype strain 10403S. Deacetylation of GlcNAc residues in PG is achieved by the *N*-deacetylase PgdA and has been shown to lead to increased lysozyme resistance (9). Since we saw a slight increase in the deacetylation of GlcNAc residues in the *eslB* mutant strain, our results indicate that the lysozyme sensitivity phenotype of the *eslB* deletion strain is independent of PgdA and that this enzyme functions properly in the mutant strain. A second enzyme required for lysozyme resistance in *L. monocytogenes* is OatA, which transfers *O*-acetyl groups to MurNAc (10, 44, 45). Using a colorimetric *O*-acetylation assay, we were able to show that PG isolated from the *eslB* mutant is less *O*-acetylated and we assume that this reduction in *O*-acetylation contributes to the lysozyme sensitivity of strain 10403SΔ*eslB*.

Growth comparisons in different media revealed that the absence of EslB results in a reduced growth in BHI broth containing high concentrations of mono- or disaccharides. One could speculate that the EslABC transporter might be a sugar transporter with a broad sugar spectrum. However, we could not identify a potential substrate binding protein encoded in the *esl* operon, which is important for substrate recognition and delivery to ABC importers. EslABC could also be involved in the export of PG components and thus affecting cell wall biosynthesis in *L. monocytogenes*. Indeed, we could show that the *eslB* mutant produces a thinner PG layer as compared to the wildtype strain, suggesting that EslABC affects PG biosynthesis. Future studies will aim to determine how the ABC transporter EslABC influences the biosynthesis and subsequent modification of PG in *L. monocytogenes*.

Absence of EslB leads to the formation of elongated cells, however, it is currently not clear how the function of EslABC is linked to cell division of *L. monocytogenes*. It seems unlikely that the activity or levels of the autolysins p60 and NamA are affected by the absence of EslB. While *iap* and *namA* mutants also form chains of cells, the cell length of individual cells is still similar to wild-type cells, however the bacteria are just unable to separate (41, 42, 46). This is in contrast to the *eslB* mutant, in which the cell length of individual cells is increased suggesting that cell division is blocked at an earlier step. In elongated cells of the *eslB* mutant, we could observe several ZapA foci, suggesting that really early cell division proteins can still be recruited in this strain. Thus, a process downstream of ZapA localization but before the construction of the actual cell septum is perturbed in the absence of EslB. EslABC could potentially affect the activity of cell division proteins or the localization of late cell division-specific proteins. Hence, deletion of *eslB* could lead to a delayed assembly of an active divisome, which could lead to an altered PG biosynthesis at the division site and an impaired cell integrity. Indeed, cells of the *eslB* mutant lysed more rapidly as compared to the *L. monocytogenes* wildtype strain 10403S when shifted from BHI broth to Tris-buffer. The autolysis of cells lacking EslB was strongly induced following growth in BHI supplemented with 0.5 M sucrose prior to the incubation in Tris-buffer. These results indicate that the *eslB* mutant is sensitive to an osmotic downshift and we hypothesize that this is due to the production of a thinner PG layer and a resulting impaired cell integrity.

Reduced lysozyme resistance is often associated with reduced virulence. An *E. faecalis* strain with a deletion in the gene coding for the peptidoglycan deacetylase PgdA, showed a reduced ability to kill *Galleria mellonella* (11). Similarly, a *S. pneumoniae pgdA* mutant showed a decreased virulence in a mouse model of infection (13). In our study, we found that inactivation of EslB does not affect the intracellular growth of *L. monocytogenes* in primary mouse macrophages or the ability to kill *Drosophila melanogaster*. These observations are consistent with a previous report that another component of the EslABC transporter, EslA, is dispensable for the ability of *L. monocytogenes* to spread from cell to cell (19). Previously, it was also shown that combined inactivation of PgdA and OatA reduced the ability of *L. monocytogenes* to grow in bone-marrow derived macrophages, whereas inactivation of PgdA alone had no impact on the virulence of *L. monocytogenes* (44). We therefore reason that the changes in PG structure and associated reduction in lysozyme resistance caused by deletion of *eslB* are not sufficient to affect the ability of *L. monocytogenes* to grow and survive in primary macrophages and flies.

Taken together, we could show that EslB is not only important for the resistance towards lysozyme, its absence also affects the autolysis, cell division and the ability of *L. monocytogenes* to grow in media containing high concentrations of sugars. Our results indicate that the ABC transporter EslABC has a direct or indirect impact on peptidoglycan biosynthesis and maintenance of cell integrity in *L. monocytogenes*.

## DATA AVAILABILITY

The Illumina reads for the *L. monocytogenes* strains 10403SΔ*eslB*_*(1)*_, 10403SΔ*eslB*_*(2)*_, 10403SΔ*eslB*_*(1)*_ compl. and 10403SΔ*eslB*_*(2)*_ compl. were deposited in the European Nucleotide Archive under the accession number PRJEB40123.

## ACKNOWLEDGEMENTS

We thank Ivan Andrew and Jaspreet Haywood from the CSC Genomics Laboratory, Hammersmith Hospital, for their help with the whole genome sequencing and Annika Gillis for help with the genome sequence analysis. We would also like to thank Charlotte S. C. Michaux and Sophie Helaine for the bone marrow-derived macrophages and Neil Singh for the support during the transmission electron microscopy experiments. We are grateful to Prof. Jörg Stülke for providing JR and LMS with laboratory space, equipment and consumables. This work was funded by the Wellcome Trust grant 210671/Z/18/Z and MRC grant MR/P011071/1 to AG, the German research foundation (DFG) grants RI 2920/1-1 and RI 2920/2-1 to JR, and the Wellcome Trust grant 207467/Z/17/Z and MRC grant MR/R00997X/1 to MSD. LMS was supported by the Göttingen Graduate School for Neurosciences, Biophysics, and Molecular Biosciences (GGNB, DFG grant GSC226/4).

