## Supplemental Material for "EslB is required for cell wall biosynthesis and modification in *Listeria monocytogenes*"

### SUPPLEMENTAL TABLES

#### Table S1: Bacterial strains used in this study

| Unique ID | Strain name and resistance | Source |
|---|---|---|
| ***Escherichia coli* strains** | | |
| ANG1264 | DH5α pKSV7; AmpR | (1) |
| 1265 | XL1-Blue pKT25; KanR | (2) |
| 1266 | XL1-Blue pKNT25; KanR | (3) |
| 1267 | XL1-Blue pUT18; AmpR | (2) |
| 1268 | XL1-Blue pUT18C; AmpR | (2) |
| 1269 | XL1-Blue pKT25-*zip*; KanR | (4) |
| 1270 | XL1-Blue pUT18C-*zip*; AmpR | (4) |
| ANG4243 | XL1-Blue pIMK3; KanR | (5) |
| ANG4236 | XL1-Blue pKSV7-Δ*eslB*; AmpR | This study |
| ANG4647 | XL1-Blue pIMK3-*eslB*; KanR | This study |
| ANG5660 | XL1-Blue pPL3e-$P_{eslA}$-*eslABC*; CamR | This study |
| ANG5661 | SM10 pPL3e-$P_{eslA}$-*eslABC*; KanR CamR | This study |
| EJR4 | XL1-Blue pKNT25-*eslA*; KanR | This study |
| EJR5 | XL1-Blue pKT25-*eslA*; KanR | This study |
| EJR6 | XL1-Blue pUT18-*eslA*; AmpR | This study |
| EJR7 | XL1-Blue pUT18C-*eslA*; AmpR | This study |
| EJR8 | XL1-Blue pKNT25-*eslB*; KanR | This study |
| EJR9 | XL1-Blue pKT25-*eslB*; KanR | This study |
| EJR10 | XL1-Blue pUT18-*eslB*; AmpR | This study |
| EJR11 | XL1-Blue pUT18C-*eslB*; AmpR | This study |
| EJR12 | XL1-Blue pKNT25-*eslC*; KanR | This study |
| EJR13 | XL1-Blue pKT25-*eslC*; KanR | This study |
| EJR14 | CLG190 pUT18-*eslC*; AmpR | This study |
| EJR15 | XL1-Blue pUT18C-*eslC*; AmpR | This study |
| EJR39 | XL1-Blue pIMK2-*mNeonGreen-zapA*; KanR | This study |
| EJR43 | XL1-Blue pKSV7-Δ*eslC*; AmpR | This study |
| EJR54 | XL1-Blue pKSV7-Δ*eslA*; AmpR | This study |
| EJR60 | S17-1 pIMK2-*mNeonGreen-zapA*; KanR | This study |
| ***Listeria monocytogenes* strains** | | |
| ANG1263 | 10403S; StrepR | (6) |
| ANG4275 | 10403SΔ$eslB_{(1)}$; StrepR | This study |
| ANG4678 | 10403S pIMK3-*eslB*; StrepR KanR | This study |
| ANG4688 | 10403SΔ$eslB_{(1)}$ pIMK3-*eslB* (or short: 10403SΔ$eslB_{(1)}$ compl.); StrepR KanR | This study |
| ANG5662 | 10403SΔ$eslB_{(2)}$; StrepR | This study |
| ANG5663 | 10403SΔ$eslB_{(2)}$ pPL3e-$P_{eslA}$-*eslABC* (or short: 10403SΔ$eslB_{(2)}$ compl.); ErmR StrepR | This study |
| LJR7 | 10403SΔ*eslC*; StrepR | This study |
| LJR21 | 10403SΔ*eslC* pPL3e-$P_{eslA}$-*eslABC* (or short: 10403SΔ*eslC* compl.); ErmR StrepR | This study |
| LJR28 | 10403S pIMK2-*mNeonGreen-zapA*; StrepR KanR | This study |
| LJR29 | 10403SΔ$eslB_{(2)}$ pIMK2-*mNeonGreen-zapA*; StrepR KanR | This study |
| LJR33 | 10403SΔ*eslA*; StrepR | This study |
| LJR34 | 10403SΔ*eslA* pPL3e-$P_{eslA}$-*eslABC* (or short: 10403SΔ*eslC* compl.); ErmR StrepR | This study |

**Table S2: Primers used in this study**

| Number | Name | Sequence |
|---|---|---|
| ANG2532 | *eslB* up fw | GCGCGGATCCCCGCAATATGTGAAATTAGTATAATG |
| ANG2533 | *eslB* up rev | TTGGAAAAGCCCTTCCCGGTACCATAACCCTCGATTAAACAT |
| ANG2534 | *eslB* down fw | TGGTACCGGGAAGGGCTTTTCCAATTGTCTAAAACGAATTAA |
| ANG2535 | *eslB* down rev | GCGCTCTAGAGCTCGCGCACTCTCATAAAC |
| ANG2812 | pIMK3-*eslB* fw | GCGCCCATGGGGTTTAATCGAGGGTTATGGTACC |
| ANG2813 | pIMK3-*eslB* rev | GCGCGTCGACTTAATTCGTTTTAGACAATTGGAAA |
| ANG3349 | pPL3e-*eslABC* fw SalI | ACGCGTCGACCTGGATGTGGCGTAAGGG |
| ANG3350 | pPL3e-*eslABC* rev BamHI | CGCGGATCCCATAACTTTGTCCCGATTGTCC |
| JR39 | Neon_rev | GCCACTACTTGTCTTATAGAGTTCATCCATACCCA |
| JR40 | Neon_ZapA_dw_f | GAACTCTATAAGACAAGTAGTGGCCTAAACGAATTT |
| JR44 | B2H EslA fw XbaI | CTAGTCTAGAAAAAATTCGGAACTTAACTAAAAAGATG |
| JR45 | B2H EslA rev KpnI | CGGGGTACCGGAATCTCGTAATAATCTATTTTATCATTTG |
| JR46 | B2H EslB fw XbaI | CTAGTCTAGAATTTAATCGAGGGTTATGGTACC |
| JR47 | B2H EslB rev BamHI | CGCGGATCCCAATTCGTTTTAGACAATTGGAAAAGC |
| JR48 | B2H EslC fw XbaI | CTAGTCTAGAAACGACAGAAACAGAAACGATTAAAC |
| JR49 | B2H EslC rev KpnI | CGGGGTACCGGTAAGGATGGAATATTTTTCTTTTTCTTC |
| JR73 | p3-mNeon-Zap fw NcoI | CATGCCATGGTTTCGAAAGGAGAGGAGGATAAT |
| JR74 | p3-mNeon-Zap rev SalI | GCGCGTCGACTTAATCTCTTCCTTTAATTCGAGC |
| LMS155 | *eslC* up fwd | CGCGGATCCAATTTTTCTTCGCCATCGCTTCC |
| LMS156 | *eslC* down rev | CGGGGTACCCTGCATCATTACCATATAAACGGACG |
| LMS157 | *eslC* down fwd | CTACATTAATATGCTTCGTTAAATTAGCTTTCAATTCAGC |
| LMS158 | *eslC* up rev | ATTTAACGAAGCATATTAATGTAGCGTTAAGTCCGATTGTTAA |
| LMS159 | *eslA* down rev | CCGGAATTCGCTTCTTTGTTTTTATACCCAGCATTAC |
| LMS160 | *eslA* up fwd | CGCGGATCCGAGTACACTTAATATTACGCTAAAAATCACAG |
| LMS161 | *eslA* up rev | AAAAGATGGACGCGATAGAAGACATTTTCACGGTGC |
| LMS162 | *eslA* down fwd | TGTCTTCTATCGCGTCCATCTTTTTAGTTAAGTTCCGAA |
|  | EGD-E_ActA_L1 | AAGCGAAGGAAGAACCAGGG |
|  | EGD-E_ActA_R1 | CCCCTAAAGAGAACACGCCA |

**Table S3: Identified sequence alterations in *L. monocytogenes eslB* deletion and complementation strains.**

| Strain number | Reference position[1] | Type[2] | Ref[3] | Allele[4] | Frequency[5] | Average Quality[6] | Annotations | AA change[7] |
|---|---|---|---|---|---|---|---|---|
| ANG4275 10403SΔ*eslB*$_{(1)}$ | 2425786-2425875 | DEL | | | 100% | | *lmo2396*, internalin | 30 aa deletion |
| ANG4688 10403SΔ*eslB*$_{(1)}$ compl. | 2366318 | SNV | C | A | 100% | 37.04 | *lmo2342*, 16S pseudo uridylate synthase | Val186Leu |
| ANG5662 10403SΔ*eslB*$_{(2)}$ | – | | | | | | | |
| ANG5663 10403SΔ*eslB*$_{(2)}$ compl. | 2058478 | DEL | A | – | 100% | 36.33 | *lmo2022,* NifS-like protein required for NAD biosynthesis | His171fs |

[1] Reference position is based on the position in the *L. monocytogenes* 10403S reference genome (NC_01744).

[2] Type of mutation: SNV = single nucleotide variant; DEL = nucleotide deletion.

[3] Ref indicates base in reference genome.

[4] Allele indicates base at the same position in the sequenced strain.

[5] Frequency at which the base change was found in the sequenced strain.

[6] Average quality score.

[7] AA change indicates the resulting amino acid change in the protein found in the reference strains as compared to the sequenced strain.

### SUPPLEMENTAL FIGURES

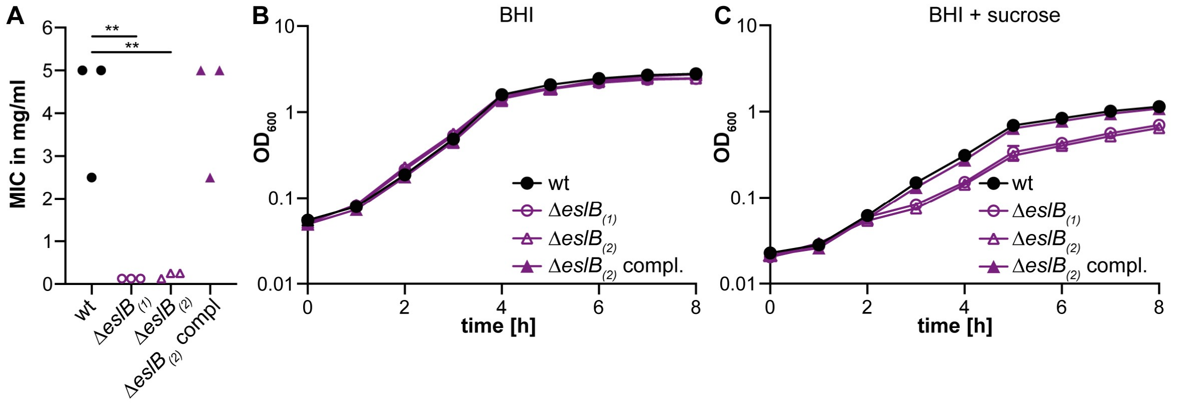


**Figure S1: Comparison of lysozyme sensitivity and growth of two different *eslB* mutants and a complementation strain.** (A) The minimal inhibitory concentration for lysozyme was determined for *L. monocytogenes* strains 10403S (wt), 10403SΔ*eslB$_{(1)}$*, 10403SΔ*eslB$_{(2)}$* and 10403SΔ*eslB$_{(2)}$* compl. using a microbroth dilution assay. The results of three independent experiments are shown. For statistical analysis, a one-way ANOVA followed by a Dunnett's multiple comparisons test was used (** $p$≤0.01). (B-C) Growth analysis of *L. monocytogenes* strains 10403S (wt), 10403SΔ*eslB$_{(1)}$*, 10403SΔ*eslB$_{(2)}$* and 10403SΔ*eslB$_{(2)}$* compl. in (B) BHI broth or (C) BHI broth containing 0.5 M sucrose. All cultures were incubated at 37°C and the bacterial growth monitored by determining $OD_{600}$ readings at hourly intervals. The average $OD_{600}$ readings and standard deviations were calculated from three independent experiments and plotted.

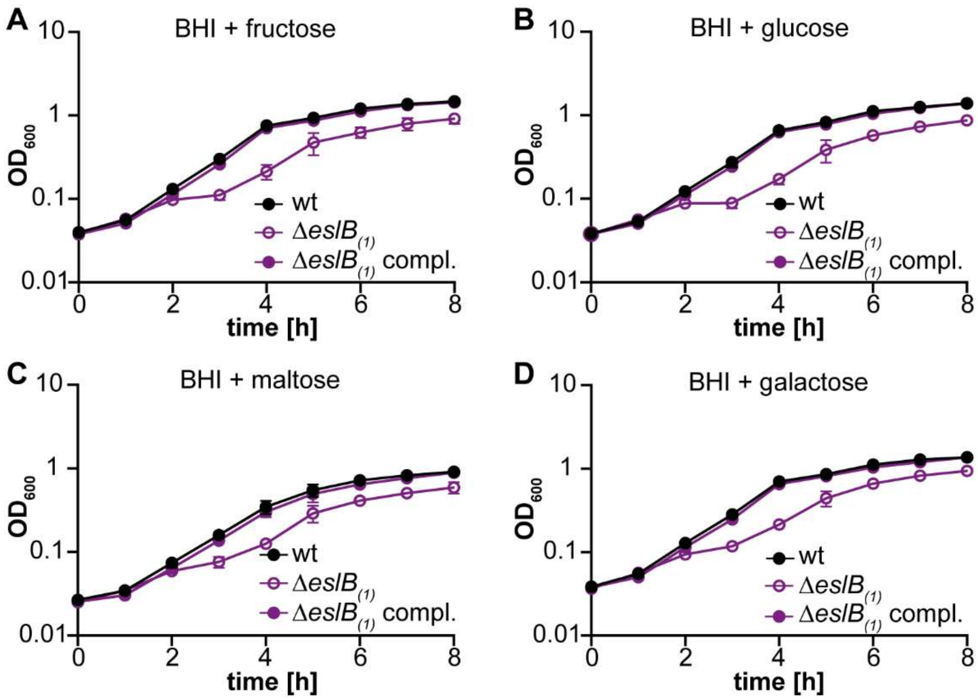


**Figure S2: The *L. monocytogenes eslB* mutant has a growth defect in BHI medium containing 0.5 M added sugar.** *L. monocytogenes* strains 10403S (wt), 10403SΔ*eslB(1)* and the complementation strain 10403SΔ*eslB(1)* compl. were grown in BHI broth containing (A) 0.5 M fructose, (B) 0.5 M glucose, (C) 0.5 M maltose or (D) 0.5 M galactose. Strain 10403SΔ*eslB(1)* compl. was grown in the presence of 1 mM IPTG. All cultures were incubated at 37°C and bacterial growth was monitored by determining $OD_{600}$ readings at hourly intervals. Average $OD_{600}$ values and standard deviations were calculated from three independent experiments and plotted. Of note, some of the error bars are too small to be seen on the graph.
